# Joint operation of the CO_2_ concentrating mechanism and photorespiration in green algae during acclimation to limiting CO_2_

**DOI:** 10.1101/2024.09.18.613642

**Authors:** Ousmane Dao, Marie Bertrand, Saleh Alseekh, Florian Veillet, Pascaline Auroy, Phuong-Chi Nguyen, Bertrand Légeret, Virginie Epting, Amélie Morin, Stephan Cuiné, Caroline Monteil, Luke C.M. Mackinder, Adrien Burlacot, Anja Krieger-Liszkay, Andreas P.M. Weber, Alisdair R. Fernie, Gilles Peltier, Yonghua Li-Beisson

## Abstract

Due to low availability of CO_2_ in aquatic environment, microalgae have evolved a CO_2_ concentrating mechanism (CCM). It has long been thought that operation of CCM would suppress photorespiration by increasing the CO_2_ concentration at the Rubisco active site, but experimental evidence is scarce. To better explore the function of photorespiration in algae, we first characterized a *Chlamydomonas reinhardtii* mutant defected in low-CO_2_ inducible 20 (LCI20) and show that LCI20 is a chloroplast-envelope glutamate/malate transporter playing a role in photorespiration. By monitoring growth and glycolate excretion in mutants deficient in either CCM or photorespiration, we conclude that: *i*. CCM induction does not depend on photorespiration, *ii*. glycolate excretion protects algal cells from the toxicity of unmetabolized photorespiratory intermediates, *iii*. photorespiration is active at low CO_2_ when the CCM is operational. This work provides a foundation for a better understanding of the carbon cycle in the ocean where significant glycolate concentrations have been found.

## Introduction

Microalgae account for nearly half of the photosynthetic CO_2_ fixation on Earth^1,2^. They contribute to climate change mitigation by assimilating CO_2_ and are promising candidates for the production of biofuels biobased compounds^3–5^. Due to the low availability of CO_2_ in water, most algal cells have developed biophysical CO_2_ concentrating mechanisms (CCMs), thus increasing the CO_2_ concentration at the catalytic site of Rubisco and favoring the carboxylase reaction at the expense of the oxygenase. Microalgal biophysical CCMs generally involve carbonic anhydrases located in different cellular compartments, bicarbonate and/or CO_2_ channels or transporters, and in most cases a specific cellular compartment the pyrenoid where Rubisco is packed in a liquid phase separated organelle and where CO_2_ is concentrated^6–10^. As a consequence of the CCM operation, the production of 2-phosphoglycolate (2-PG) resulting from the oxygenase activity of Rubisco is reduced^11,12^. Because of its inhibitory effect on the Calvin cycle, 2-PG is metabolized and recycled through photorespiration. Photorespiration consists of multiple metabolic reactions distributed across different subcellular compartments thus requiring inter-organelle communication^13^. During photorespiration, the 2-PG is converted in the chloroplast into glycolate by the 2-PG phosphatase PGP1. In the model green microalga *Chlamydomonas reinhardtii* (hereafter *Chlamydomonas*), part of the glycolate is transferred to mitochondria and further metabolized into glyoxylate, glycine and serine, the other part is excreted out of the cell^14–16^. During acclimation to low CO_2_, CCM and photorespiratory genes, which share the same master regulator CIA5, are simultaneously induced^17–19^. Such a feature is intriguing since the activity of CCM supposedly inhibits photorespiration. It was proposed that photorespiration is transiently operational during the acclimation from high to low CO_2_ conditions until the CCM is fully induced^7,12^, photorespiratory metabolites such as 2-PG possibly acting as signaling molecules triggering CCM induction^12,20–23^.

Two types of CCM regimes have been described in *Chlamydomonas* depending on external CO_2_ levels, the low-CO_2_-based CCM (CO_2_ air level), in which cells preferentially transport CO_2_, and the very-low-CO_2_-based CCM, in which specific transporters are induced, such as the ATP-binding cassette transporter High Light Activated 3 (HLA3) involved in active bicarbonate uptake^24,25^.7 During CCM induction, mitochondria migrate from the chloroplast cup to the periphery of the cell at the vicinity of the plasma membrane^26^. The contribution of mitochondria to the energy supply of the algal CCM has been recently evidenced, and energy trafficking between chloroplast and mitochondria was suggested to supply ATP to CCM bicarbonate transporters^27,28^. Actually, energy exchange between subcellular organelles was evidenced decades ago in microalgae^30–33^ and has been re-examined more recently^29,34–36^. The exchange of reducing power between organelles may operate via “malate shuttles” composed of malate dehydrogenases (MDHs) and membrane transporters of dicarboxylic acids^37–40^. Despite the existence of several genes encoding putative chloroplast malate transporters^40–42^, none of them has been characterized in *Chlamydomonas*. Based on the increased expression of low-CO_2_ inducible 20 (*LCI20*) gene, which encodes a putative malate/2-oxoglutarate transporter, during acclimation to limiting CO_2_^17–19^, it was proposed that LCI20 may be involved as a malate shuttle in the energy trafficking between chloroplast and mitochondria to feed CCM bicarbonate transporters^43^. By another way, photorespiration may also contribute to the energy supply to external CCM transporters^43,44^. Indeed, during photorespiration NADH is produced during the mitochondrial condensation of two glycine molecules into one serine, the NADH being converted into ATP by the mitochondrial respiratory chain. Until now, the nature of metabolic pathway and the identity of proteins involved in the energy network transferring photosynthetic energy from the chloroplast towards mitochondria to feed the external CCM transporters remain largely unexplored.

In this study, we aimed at better characterizing the relationship between CCM and photorespiration in green algae. For this purpose, we characterized a *Chlamydomonas* mutant deficient in LCI20 and show that LCI20 is a malate/glutamate transporter involved in photorespiration by supplying amino groups for the mitochondrial conversion of glyoxylate into glycine. By comparing growth properties and glycolate excretion in mutants affected in photorespiration or CCM under various photorespiration regimes, we conclude that photorespiratory metabolites do not contribute to CCM induction, that glycolate excretion avoids toxicity of non-metabolized photorespiratory intermediates and that downstream glycolate metabolism, although occurring when the CCM is functioning under very low CO_2_, is not required for CCM operation.

## Results

### *LCI20* encodes a putative chloroplast malate transporter whose expression is modulated by light and CO_2_ levels

Three putative chloroplast 2-oxoglutarate/malate transporters (annotated as OMT1, OMT2 and LCI20) orthologues of previously characterized Arabidopsis DiT1 and DiT2 malate transporters were identified in the Chlamydomonas genome^41,42^. Previous transcriptomic analysis performed during a day-night cycle or during limiting CO_2_ acclimation showed that the expression of *LCI20* (but not that of *OMT1* and *OMT2*) is strongly induced at the beginning of the light period^45^ or after the transition from H-CO_2_ (2%) to L-CO_2_ (0.04%) or VL-CO_2_ (0.01%)^17–19^ (**Supplementary Fig. 1a, b**). A phylogenetic analysis revealed that LCI20, which belongs to the GreenCut2^46^, is widespread in *Chlorophyta*, closely related to DiT2 transporters, and evolutionarily divergent from OMT family transporters (**Supplementary Fig. 1c and 2**). Furthermore, LCI20 has recently been localized in the chloroplast but the enrichment in the envelope was not clearly shown^47,48^. Here, we reexamined the strain expressing *LCI20*-mVenus under the control of *PSAD* promoter and we can see clearly that LCI20 is localized to the chloroplast envelope (**Fig. 1a and Supplementary Fig. 3**).

**Fig. 1.**
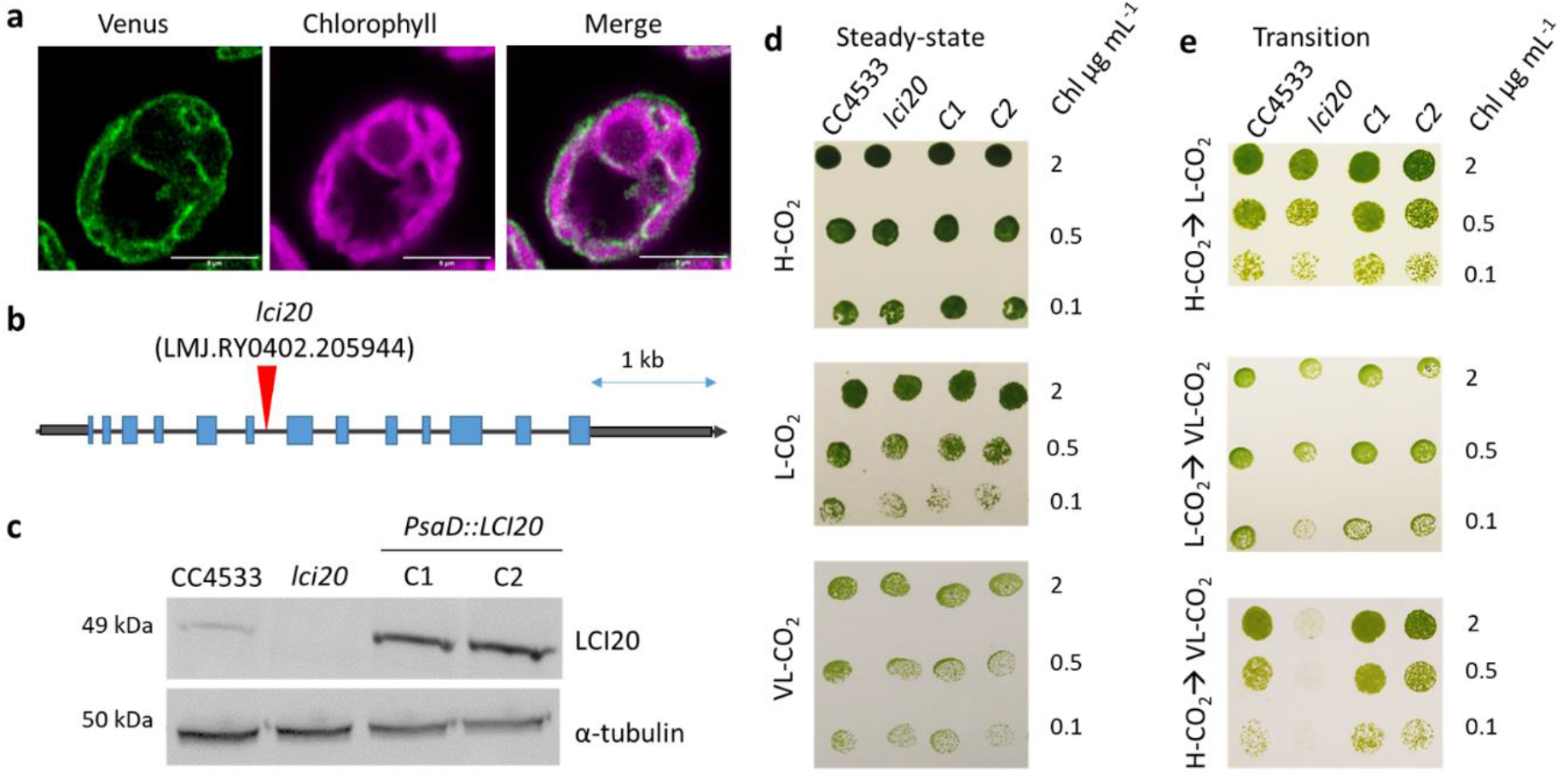
LCI20 subcellular localization, characterization and complementation of a *lci20* insertional mutant. (**a**) Subcellular localization of LCI20 protein fused with the Venus fluorescent reporter. False colours were used to represent Venus (green) and chlorophyll (magenta) fluorescence signals. Scale bar, 5 µm. (**b**) Genomic structure of *LCI20* gene and insertion site of the paromomycin resistance cassette in the CLiP *lci20* mutant. Gray boxes at both extremities represent the 5’ and 3’UTR, respectively. Exons are colored blue, and the position of cassette insertion is indicated by red arrows. (**c**) Immunoblot analysis using anti-LCI20 antibodies. Uncropped immunoblots are provided as a Source Data file. (**d, e**) Photoautotrophic growth of *lci20*, WT and complemented lines (C1, C2) on agar plates exposed to various CO_2_ regimes. Cells were grown photo-autotrophically in liquid culture under air CO_2_ level (**d**), or air supplemented with 2% CO_2_ (**e**) prior to spot testing. Images were taken 3 days (H-CO_2_ and LCO_2_) or 5 days (VL-CO_2_) after growing under 80 μmol photons m^−2^ s^−1^.

### The growth of *lci20* is impaired during a transition from high to very low CO2

A mutant harboring an insertion in the sixth intron of the *LCI20* locus was obtained from the Chlamydomonas library project (CLiP) collection^49^ and we named the mutant here as *lci20* (**Fig. 1b**). We confirmed the insertion of the paromomycin resistance cassette and the absence of the *LCI20* transcript in *lci20* (**Supplementary Fig. 4a, b**). We then performed genetic complementation of *lci20* by expressing the full-length genomic *LCI20* coding sequence driven by the PSAD promoter (**Supplementary Fig. 4a, b**). A LCI20 antibody was produced in this study and used to show the absence of LCI20 in the mutant and its presence in WT and the complemented lines (**Fig. 1*C*)**.

The growth of *lci20* was then investigated under photoautotrophic conditions in both solid and liquid cultures under various CO_2_ levels (**Fig. 1d, e; Supplementary Fig. 4c, d**). On solid media, *lci20* grew normally under H-CO_2_, L-CO_2_ or VL-CO_2_ when cells were previously acclimated to these conditions (**Fig. 1d**). However, the growth of *lci20* was severely impaired when transitioning from H-CO_2_ to VL-CO_2_ (**Fig. 1e**), with smaller decreases in growth observed after a transition from H-CO_2_ to L-CO_2_ or from L-CO_2_ to VL-CO_2_ (**Fig. 1e)**. In liquid media, while *lci20* grew normally in H-CO_2_ or during transition from H-CO_2_ to L-CO_2_, its growth was reduced when expressed on the basis of cell volume during transition from H-CO_2_ to VL-CO_2_ **(Supplementary Fig. 4c, d)**. Based on these results, we conclude that LCI20 is required for photoautotrophic growth when cells are subjected to a sudden and severe CO_2_ limitation.

### *lci20* is affected in photorespiratory glycolate metabolism

To determine whether the mutant phenotype is due to a role of LCI20 in the induction of the CCM or in photorespiration, we performed growth assays on agar plates by transferring H-CO_2_ grown cells into VL-CO_2_ under photorespiratory (21% O_2_) or non-photorespiratory (2% O_2_) conditions (**Fig. 2a**). As a control, we used the CCM1 transcription factor knockout mutant *cia5*, which is defective in the transcriptional induction of CCM and of the photorespiratory pathway^11,44,50,51^. *cia5* grew normally under H-CO_2_ but growth was completely abolished when cells were transferred to VL-CO_2_, irrespective of O_2_ levels (**Fig. 2a**), thus indicating that a defect in the CCM, but not in photorespiration, prevented growth of the *cia5* mutant under VL-CO_2_. Note that similar growth was observed when wild-type cells were exposed to VL-CO_2_ either under photorespiratory conditions (21% O_2_) or non-photorespiratory conditions (2% O_2_), indicating that photorespiration is not required for CCM induction. In contrast, *lci20* grew poorly after transition under photorespiratory conditions (VL-CO_2_; 21% O_2_), but the growth was unaffected under non-photorespiratory conditions (VL-CO_2_; 2% O_2_) (**Fig. 2a**), indicating that *lci20* is defected in photorespiration rather than in CCM.

**Fig. 2.**
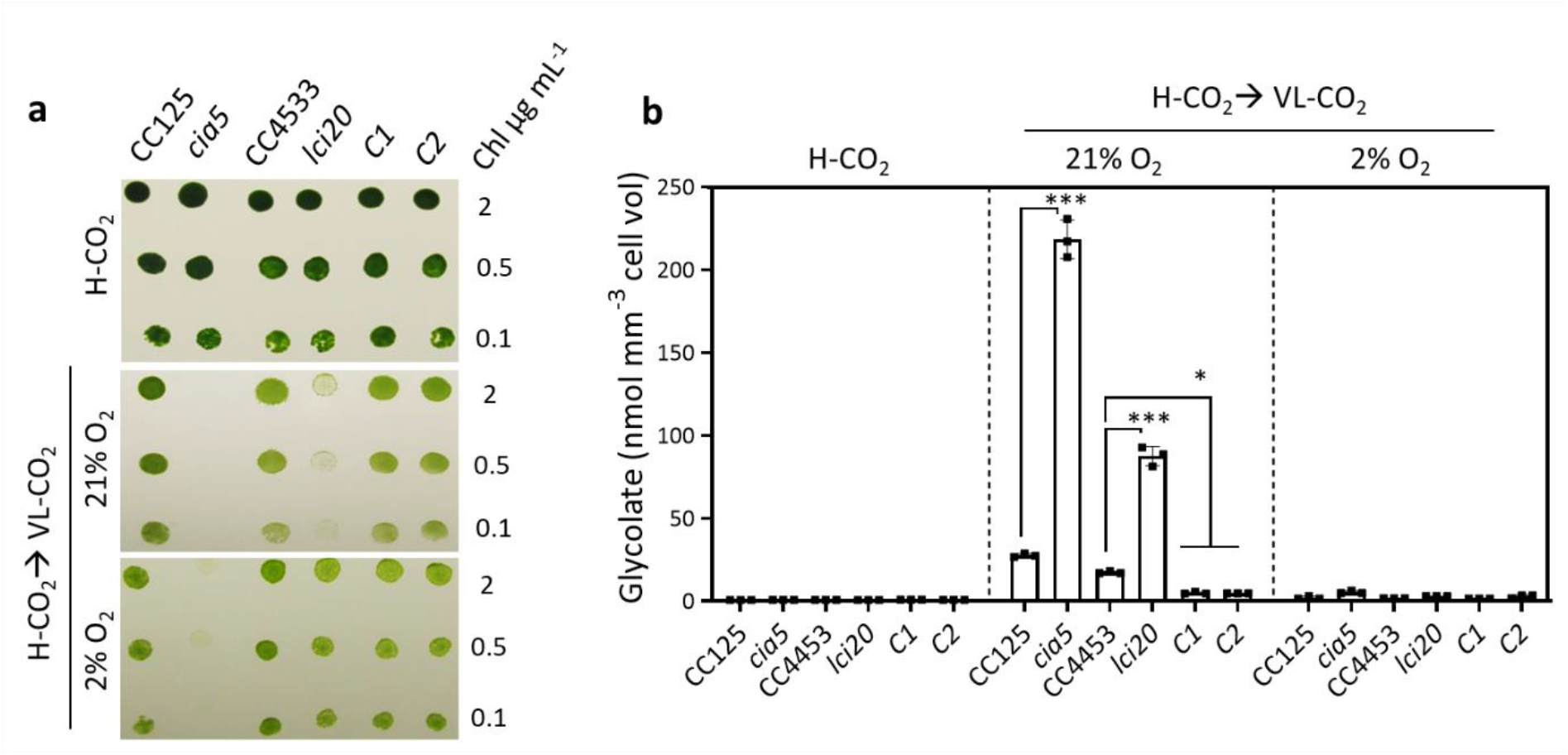
*lci20* is affected in the photorespiratory glycolate metabolism. (**a**) Growth performance of *cia5, lci20* mutants and their respective controls when transferring photo-autotrophically grown cells in liquid cultures under H-CO_2_ to agar plates exposed to VL-CO_2_ either under 21% or 2% O_2_ under 80 μmol photons m^−2^ s^−1^. Images were taken after 3 days (H-CO_2_) or 5 days (VL-CO_2_) of growth. (**b**) Quantification of the glycolate concentration in the liquid medium after 20 h growth of cultures either under H-CO_2_ or upon transition for H-CO_2_ to VL-CO_2_ either in the presence of 21% or 2% O_2_. Bars show the average and dots show data from independent biological replicates (n=3 ± SD). Asterisks represent statistically significant difference compared to the wild-type strains (* *p* ≤ 0.05, ** *p* ≤ 0.01, *** *p* ≤ 0.001 and **** *p* ≤ 0.0001) using one-way ANOVA.

Glycolate is excreted into the culture medium by *Chlamydomonas* cells when either the CCM or the glycolate metabolism is defective^14,16,44,52,53^. The glycolate concentration of the culture medium was measured after 20 h acclimation to VL-CO_2_, both *cia5* and *lci20* mutants showing a significantly higher excretion compared to their respective controls (**Fig. 2b**). Note that we did not detect glycolate excretion under H-CO_2_ or VL-CO_2_ at 2% O_2_ conditions when photorespiration is strongly reduced (**Fig. 2b**). We conclude that *lci20* is defective in the photorespiratory glycolate metabolism, which is triggered when *Chlamydomonas* cells are transferred from H-CO_2_ to VL-CO_2_.

### Changes in abundance of photorespiration and CCM proteins during cells’ response to different CO_2_ and O_2_ levels

To identify possible causes for the growth defect and glycolate over-excretion observed in *lci20*, we probed the abundance of different photorespiratory enzymes including the glycolate dehydrogenase (GYD1), hydroxypyruvate reductase (HPR1) and glycine cleavage system P (GCSP) proteins together with CCM-related proteins HLA3, low CO_2_-inducible 1 (LCI1) and low CO_2_-inducible C (LCIC) following a transfer from H-CO_2_ to VL-CO_2_ either at 21% or at 2% O_2_ (**Fig. 3a**). We observed a lower accumulation of GYD1 in *lci20* compared to control strains when the transfer was performed under photorespiratory conditions (21% O_2_), but no decrease under non-photorespiratory conditions (2% O_2_) (**Fig. 3a**). The abundances of other photorespiration or CCM enzymes were similar in *lci20* and control strains (**Fig. 3a**), with the exception of LCIC which showed a slightly lower abundance in *lci20* under VL-CO_2_ independent of photorespiration (**Fig. 3a**).

**Fig. 3.**
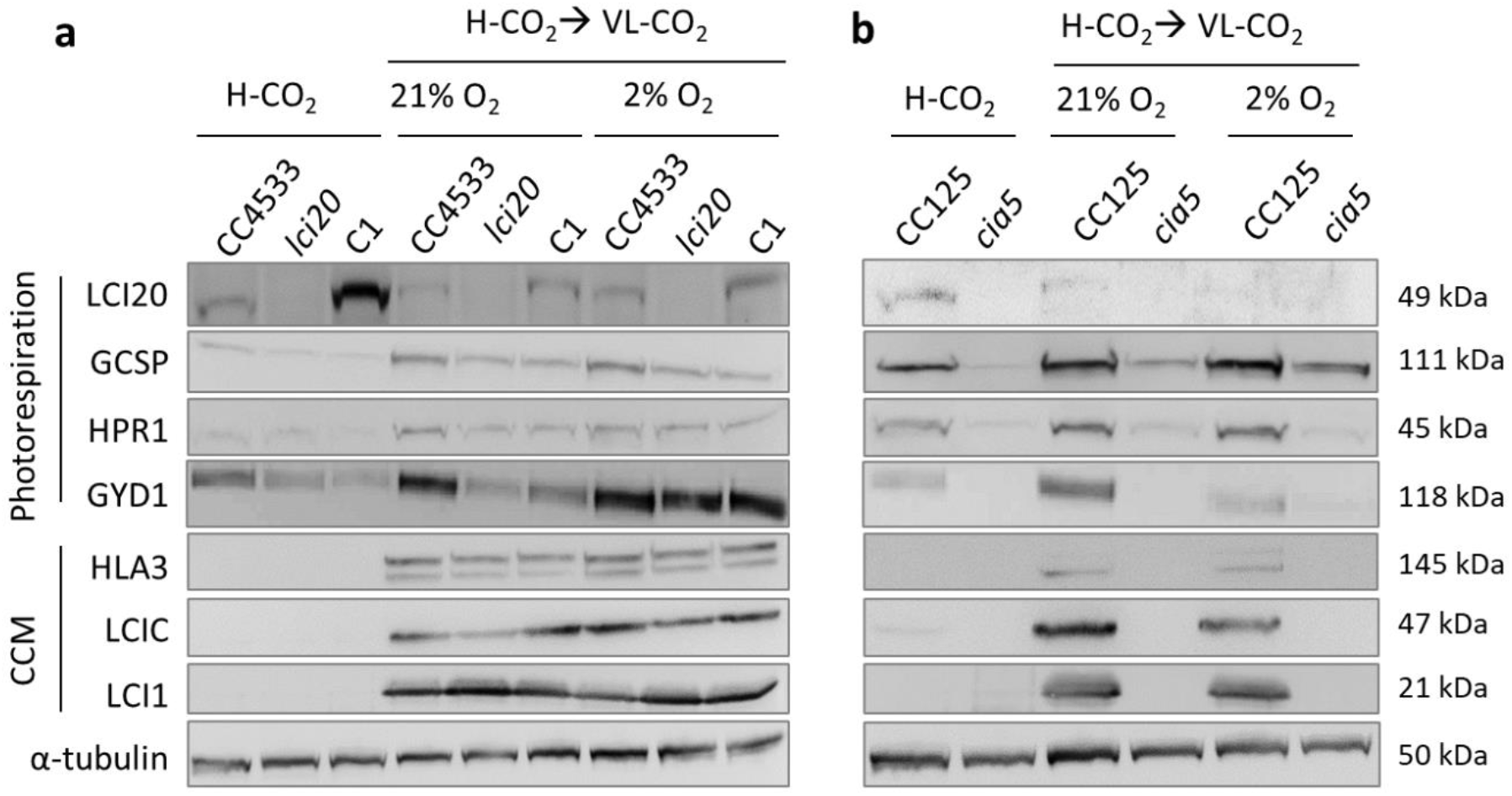
Effect of CO_2_ and O_2_ concentrations on the abundance of photorespiration and CCM proteins. (**a**) Immunoblot analysis of representative CCM and photorespiration related proteins in *lci20*, its wild-type control CC4533 and one complemented line (C1). (**b**) Immunoblot analysis of representative CCM and photorespiration related proteins in *cia5* and its wild-type control CC125. α-tubulin was used as a loading control. Cells were cultivated photo-autotrophically under H-CO_2_ and 80 μmol photons m^−2^ s^−1^ and then acclimated for 20 h at the indicated CO_2_ and O_2_ levels prior to immunoblot analysis. Uncropped immunoblots are provided as a Source Data file.

In *Chlamydomonas*, photorespiration and CCM genes are co-regulated during acclimation to sub-optimal CO_2_ and controlled by a common transcription factor CIA5. We then monitored the accumulation of photorespiratory enzymes (GYD1, HPR1 and GCSP) and of CCM-related proteins (HLA3, LCI1 and LCIC) following acclimation to different CO_2_ and O_2_ levels in *cia5* and its wild-type control (**Fig. 3b**). Whereas CCM and photorespiratory proteins were induced after 20 h of acclimation to VL-CO_2_ in a CIA5-dependent manner, only photorespiratory proteins were detected under H-CO_2_ (**Fig. 3b**), indicating that photorespiratory enzymes and CCM proteins are regulated differently at the protein level.

Then, to better understand the reasons of the growth defect observed in *lci20*, and further determine whether it may be caused by a decrease of GYD1 abundance, we have isolated and characterized two CLiP mutants of *Chlamydomonas* harboring insertions in introns of the *GYD1* gene (**Fig. 4a**) and validated to be deficient in the GYD1 protein by immunoblot (**Fig. 4b**). Both mutants showed higher glycolate excretion than the control strain (**Fig. 4c**). However, the growth of both *gyd1* mutants was similar to the control wild-type strain after a transition from H-CO_2_ to VL-CO_2_ regardless of O_2_ levels (**Fig. 4d**). It turns out that *gyd1* mutants do not require H-CO_2_ to grow, which is in contrast to a previous work where the H-CO_2_ requiring 89 mutant (HCR89) is reported to harbor a mutation at the *GYD1* locus^44^. Therefore, the growth defect of *lci20* observed during H-CO_2_ to VL-CO_2_ is likely not due to a decrease in GYD1 abundance, but rather results from the accumulation of photorespiratory metabolites downstream the GYD1 reaction step. Indeed, the *gyd1* mutants excreted twice more glycolate than *lci20* mutants (**Fig. 2b and 4c**), which might explain why the *gyd1* mutants grow better than *lci20* during the transition from H-CO_2_ to VL-CO_2_.

**Fig. 4.**
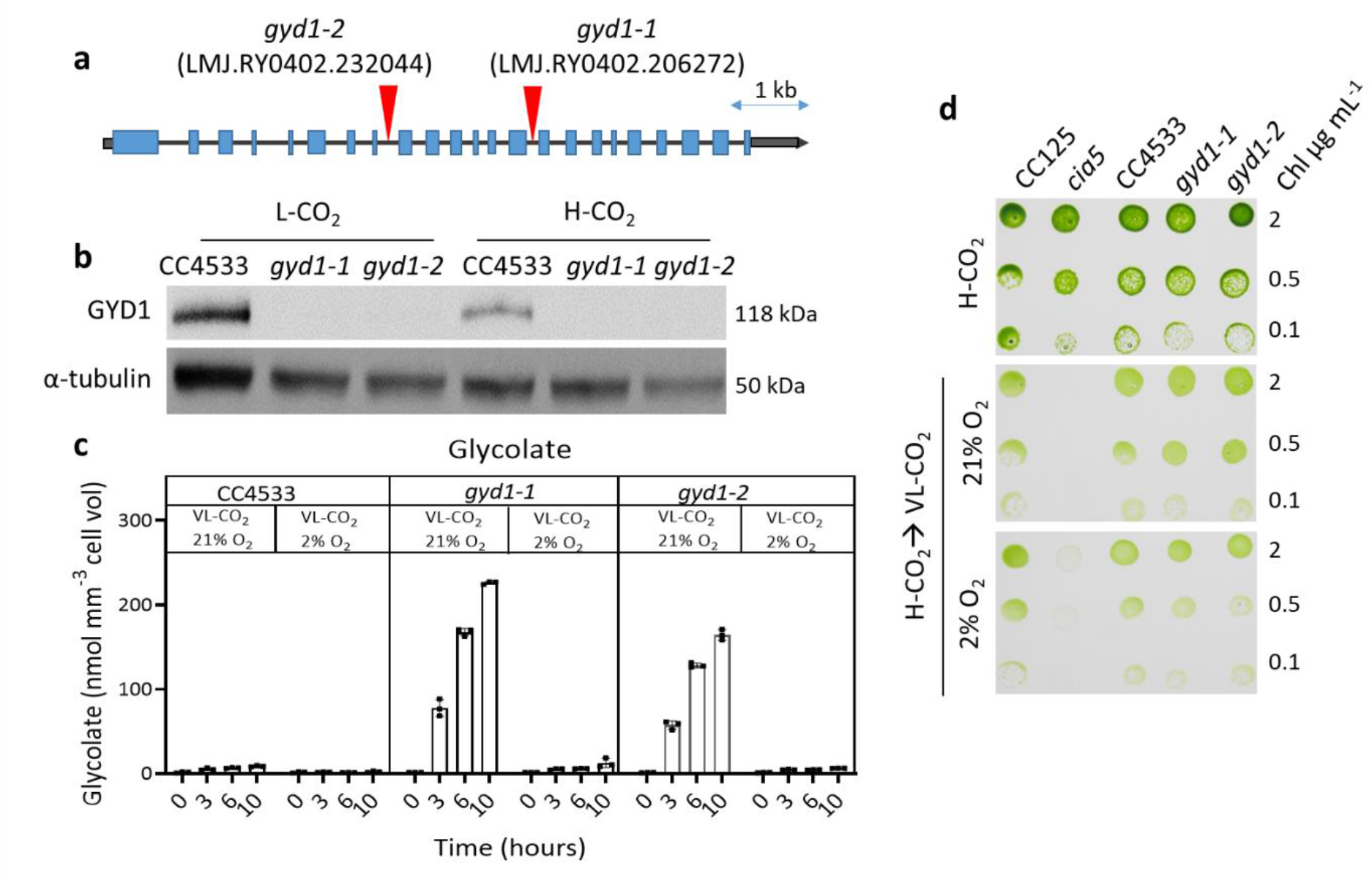
Characterization of two insertional mutants of the glycolate dehydrogenase gene *GYD1*. (**a**) Genomic structure of *GYD1* gene and insertion sites of the paromomycin resistance cassettes in two CLiP mutants *gyd1-1* and *gyd1-2*. Gray boxes at both extremities represent the 5’ and 3’UTR, respectively. Exons are colored blue and the position of cassette insertion is indicated by red arrows. (**b**) Immunoblot analysis using anti-GYD1 antibodies performed on both *gyd1* mutants and their WT control grown under H-CO_2_ or VL-CO_2_. α-tubulin was used as a loading control. Uncropped immunoblots are provided as a Source Data file. (**c**) Quantification of the glycolate concentration in the culture medium after a transition from H-CO_2_ to VL-CO_2_ at 21% or 2% O_2_. Bars show the average and dots show data from independent biological replicates (n=3 ± SD). (**d**) Photoautotrophic growth of both *gyd1* mutants, their wild-type control CC4533 and *cia5* and its wild-type control CC125 on agar plates exposed to various CO_2_ and O_2_ concentrations. Cells were grown in liquid culture in flasks photo-autotrophically under H-CO_2_ prior to spot test. Images were taken after 3 days (H-CO_2_) or 5 days (VL-CO_2_) of growth under 80 μmol photons m^−2^ s^−1^.

### Photosynthesis of *lci20* is affected under photorespiratory conditions

To investigate the consequence of impaired photorespiration on the photosynthetic capacity of *lci20*, we performed room temperature chlorophyll fluorescence measurements to assess the PSII quantum yield and PQ redox state (1-qL) in H-CO_2_ grown cells and following their acclimation to VL-CO_2_ (**Fig. 5, Supplementary Fig. 5**). Under H-CO_2_, *lci20* showed a slightly higher effective PSII quantum yield and lower PQ redox state compared to wild-type but was mainly similar to the complemented line **(Fig. 5a, d)**. Following VL-CO_2_ acclimation under photorespiratory conditions (21% O_2_), *lci20* showed a significantly reduced effective PSII quantum yield and elevated PQ redox state compared to the control strains (wild-type and complemented line) (**Fig. 5b, e**). These effects were suppressed under non-photorespiratory conditions (VL-CO_2_, 2% O_2_) (**Fig. 5c, f**), thus indicating that the accumulation of non-metabolized photorespiratory compounds may be responsible for the inhibition.

**Fig. 5.**
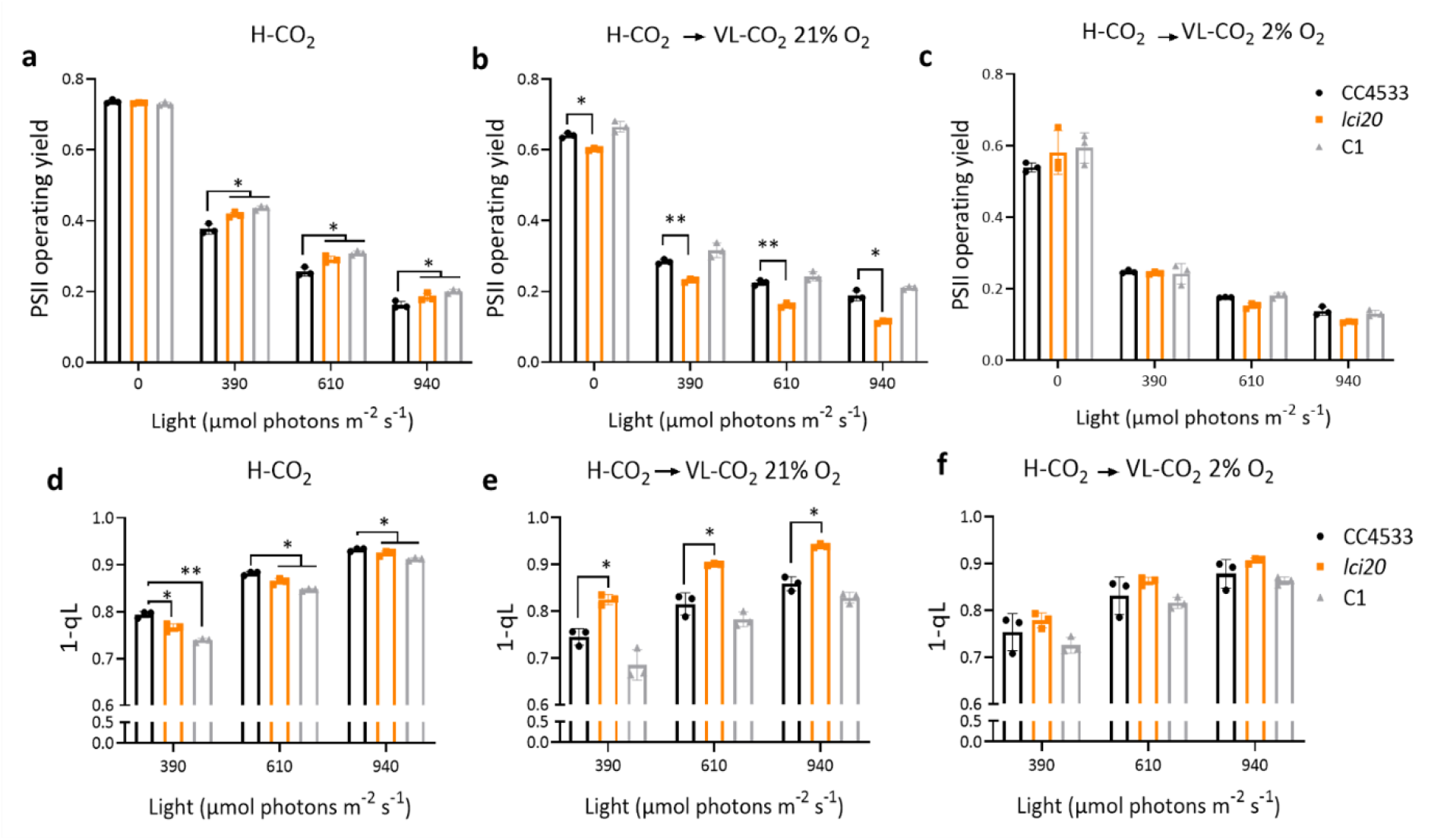
Photosynthetic activity measured by chlorophyll fluorescence is reduced in *lci20* during acclimation to VL-CO_2_ under photorespiratory (21% O_2_) but not under non-photorespiratory (2% O_2_) conditions. (**a-c**) PSII operating yields measured under actinic light in cells grown photo-autotrophically (**a**) under H-CO_2_, (**b**) during acclimation from H-CO_2_ to VL-CO_2_ at 21% O_2_ or (**c**) at 2% O_2_ for 20 h. (**d-f**) PQ redox state measured as the (1-qL) parameter in the same samples as in (**a-c**). Cells were exposed for 1 min at the indicated light intensity. Bars show the average and dots show data from independent biological replicates (n=3 ± SD). Asterisks represent statistically significant difference compared to the wild-type strain CC4533 (* *p* ≤ 0.05, ** *p* ≤ 0.01, *** *p* ≤ 0.001 and **** *p* ≤ 0.0001) using one-way ANOVA.

### Metabolomic analyses reveal an increased accumulation of malate and glutamate in *lci20*

To investigate the metabolic changes occurring in *lci20* and obtain clues to the metabolites transported by LCI20, we performed a metabolomic analysis in the different strains grown under photoautotrophic H-CO_2_, and during transitions from H-CO_2_ to L-CO_2_ and from H-CO_2_ to VL-CO_2_ (**Fig. 6; Supplementary Fig. 6; Supplementary Data 1**). The metabolite profile of H-CO_2_-grown *lci20* cells was comparable to that of control strains (**Fig. 6a; Supplementary Fig. 6; Supplementary Data 1**). After transition from H-CO_2_ to L-CO_2_ or to VL-CO_2_, we observed a higher accumulation of malic acid and glutamic acid in *lci20* when cells were cultivated under photorespiratory condition (21% O_2_), the effect being suppressed under non-photorespiratory conditions (2% O_2_) (**Fig. 6b, d**). Accumulation of almost all the other metabolites detected was not significantly affected in *lci20* under the conditions tested (**Supplementary Fig. 6; Supplementary Data 1**). Based on these results, we propose that LCI20 is involved in the transport of malate and glutamate during photorespiration and that disruption of glutamate transport impairs photorespiratory glycolate metabolism at the step of glyoxylate conversion to glycine thus explaining the *lci20* phenotypes observed during acclimation to VL-CO_2_.

**Fig. 6.**
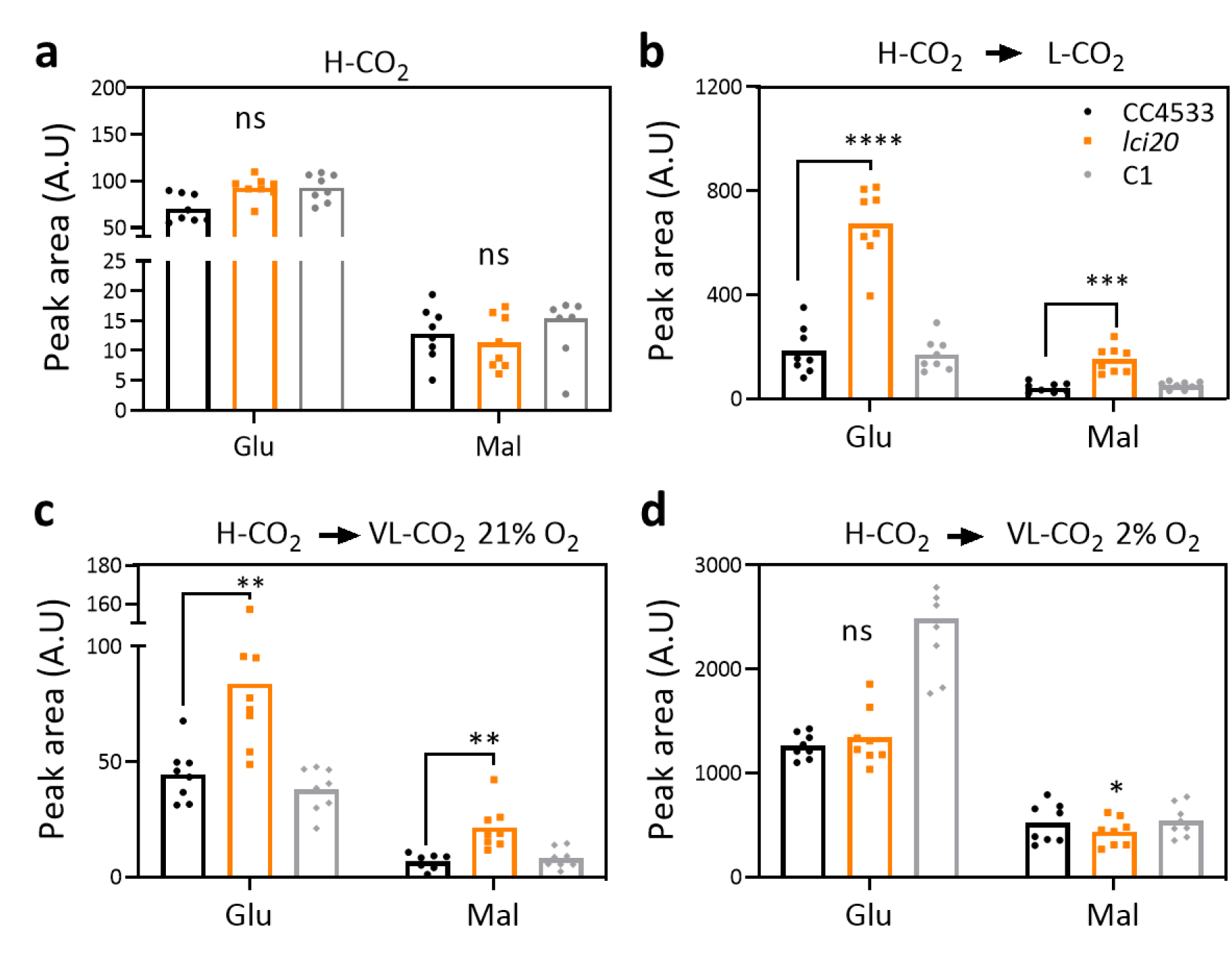
Metabolomic analysis shows increased accumulation of malate and glutamate in *lci20* during a transition from H-CO_2_ to VL-CO_2_ under photorespiratory but not under non-photorespiratory conditions. (**a-d**) Accumulation of intracellular glutamic acid and malic acid in cells grown (**a**) under H-CO_2,_ (**b**) or after transition from H-CO_2_ to L-CO_2_ for 20 h, (**c**) after transition from H-CO_2_ to VL-CO_2_ under 21% O_2_ or (**d**) under 2% O_2_ for 20 h. Bars show the average and dots show data from independent biological replicates (n=8 ± SD). Asterisks represent statistically significant difference compared to the wild-type strain CC4533 (* *p* ≤ 0.05,** *p* ≤ 0.01 and *** *p* ≤ 0.001) using one-way ANOVA.

## Discussion

In this study, we demonstrate the role of LCI20 in the photorespiratory glycolate metabolism in *Chlamydomonas* (**Fig. 7**), as evidenced by a high glycolate excretion and the marked growth defect of the *lci20* mutant during a transition from H-CO_2_ to VL-CO_2_ at ambient O_2_ levels (21%), these effects being suppressed at 2% O_2_ (**Fig. 1-2**). Many photorespiratory mutants have been previously reported in land plants and only very few in algae. In *Arabidopsis*, a defect in genes encoding photorespiratory enzymes generate a high-CO_2_ requiring phenotype, which is attributed to the accumulation of non-metabolized photorespiratory intermediates^13^. In *Chlamydomonas*, the situation is not that clear, and this is further complicated by the occurrence of CCM in algae. A phosphoglycolate phosphatase 1 (*pgp1*) mutant, impaired in the conversion of 2-PG to glycolate, exhibits a growth defect under stationary conditions of limiting CO_2_ levels^22^. The high-CO_2_ growth requirement of *pgp1* was attributed to the inhibitory effect of non-metabolized 2-PG accumulating upon ribulose bisphosphate oxygenation^22^. After the conversion of 2-PG into glycolate by PGP1, glycolate can follow two different fates in *Chlamydomonas*, it can be either excreted out of the cell or metabolized into glyoxylate by GYD1 in the mitochondria. A *Chlamydomonas gyd1* mutant isolated from a H-CO_2_ requiring screening showed a high glycolate excretion during transition from H-CO_2_ to L-CO_2_^44^. However, the H-CO_2_ requiring phenotype of this mutant was not clearly evidenced. We have shown here that impairing photorespiration at the level of GYD1 does not lead to an H-CO_2_requirement but instead to a high glycolate excretion under VL-CO_2_ (**Fig. 4**). It was recently reported that growth of *Chlamydomonas hpr1* mutant is slightly impaired during the transition from H-CO_2_ to L-CO_2_, accompanied by a high glycolate excretion^55^. The slight growth defect of the *hpr1* mutant being interpreted as a consequence of glycolate hyper-excretion^55^. However, the *gyd1* mutants characterized here show high glycolate excretion rates but no growth defect in any CO_2_ conditions tested (**Fig. 4**). We therefore conclude that inhibition of photorespiration at different steps of the pathway can lead to contrasting phenotypes. While inhibition of glycolate conversion does not impair growth (*i*.*e*. in *gyd1*), the accumulation of intermediate metabolites such as 2-PG (*i*.*e*. in *pgp1*) or downstream glycolate conversion (*i*.*e*. in *hpr1*) is deleterious. Consequently, glycolate excretion could be seen as a protective mechanism preventing the toxic accumulation of photorespiratory intermediates during acclimatization to VL-CO_2_, until CCM components and photorespiratory enzymes are fully induced, thus allowing photorespiratory metabolism to be tuned to the residual Rubisco oxygenase activity which remains at VL-CO_2_ despite the presence of CCM.

**Fig. 7.**
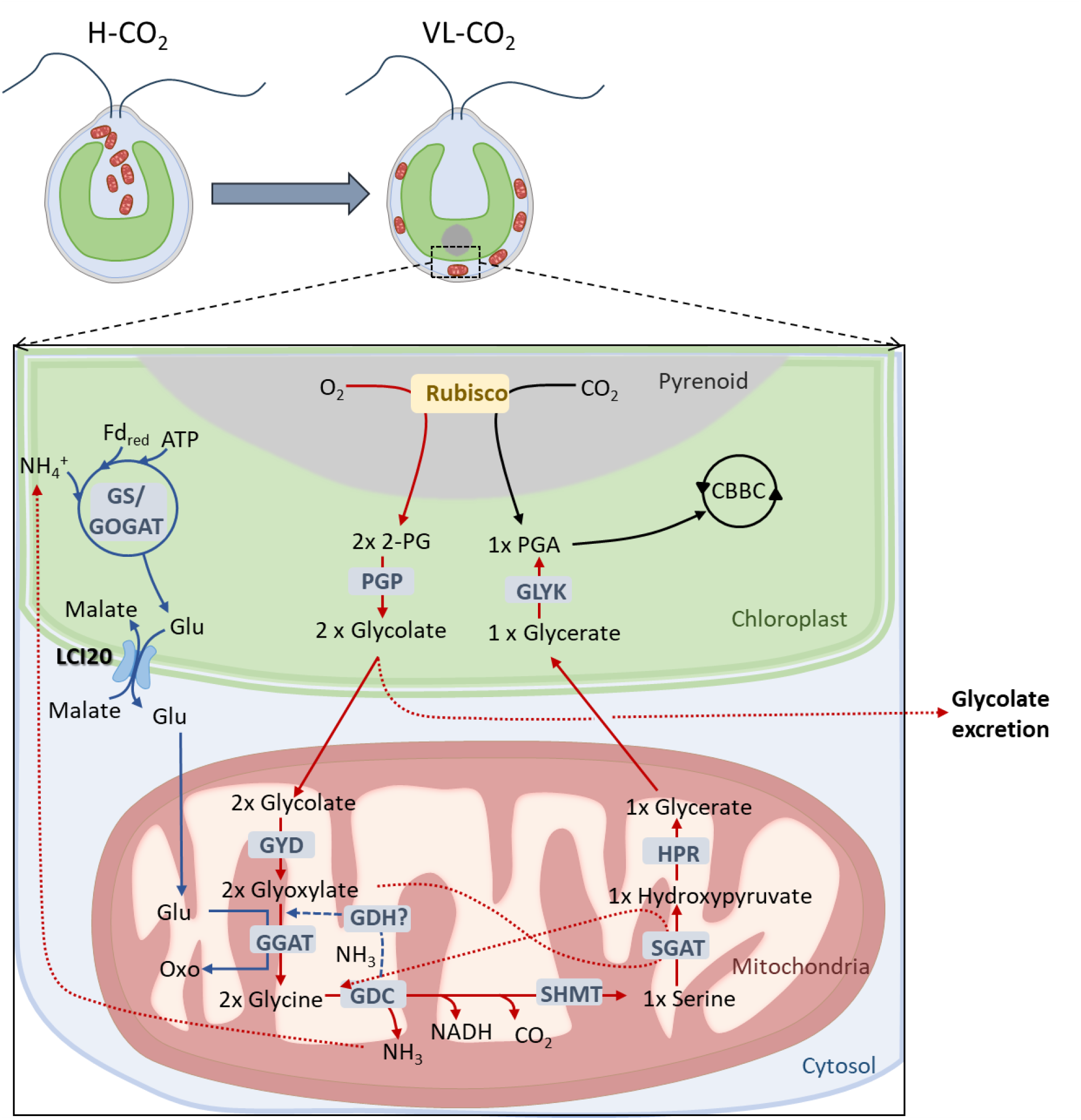
Hypothetical scheme describing the role of LCI20 during photorespiration in *Chlamydomonas*. Photorespiration is initiated by the oxygenase activity of Rubisco producing phosphoglycolate (2-PG), which is converted into glycolate by a phospho-glycolate phosphatase (PGP), and then glyoxylate by the glycolate dehydrogenase (GYD). Glyoxylate is transaminated into glycine by a glutamate-glyoxylate aminotransferase (GGAT) using glutamate as a -NH_2_ donor. During the condensation of two molecules of glycine into one serine by the glycine decarboxylase (GDC), CO_2_ and NH_3_ are released. Photorespiratory NH_3_ is likely reassimilated by the GS/COGAT as proposed in *Arabidopsis*^57,81^ and *Chlamydomonas*^15,64^. LCI20 would function as a glutamate transporter exporting glutamate out of the chloroplast in counter-exchange with malate, thus supplying a -NH_2_ donor for glyoxylate conversion into glycine. The growth phenotype of *lci20* observed during a transition from H-CO_2_ to VL-CO_2_ would result from the accumulation of glyoxylate. The absence of *lci20* phenotype under steady state L-CO_2_ or VL-CO_2_ would result from the induction of an alternative recycling pathway for the photorespiratory NH_3_, for instance the mitochondrial glutamate dehydrogenase (GDH). Abbreviations: CBBC, Calvin-Benson-Bassham cycle; Fdred, reduced ferredoxin; Glu, glutamate; GLYK, glycerate kinase; GS/GOGAT, glutamine synthetase/glutamine:oxoglutarate aminotransferase; HPR, hydroxypyruvate reductase; LCI20, low-CO2 inducible 20; Oxo, oxoglutarate; SHMT, serine hydroxy methyltransferase; SGAT, serine-glyoxylate aminotransferase; TCA, tricarboxylic acid cycle.

Unlike the above-mentioned proteins (PGP1, GYD1 and HPR1), which are all involved in the core photorespiratory pathways, LCI20 turns out to be an auxiliary protein involved in the supply of amino (-NH_2_) donor as glutamate for the conversion of glyoxylate into glycine (**Fig. 7**). In *Arabidopsis* chloroplasts, the export of glutamate in counter-exchange with malate is mediated by the dicarboxylate transporter DiT2.1 that is required for photorespiratory nitrogen assimilation and displays a 60% homology with the *Chlamydomonas* LCI20^56–59^. Based on our metabolomic analysis, we suggest that LCI20 mediates the export of glutamate in counter exchange with malate across the chloroplast inner envelope (**Fig. 6; Supplementary Fig. 6**). Impairing glutamate export by knocking out LCI20 likely disrupts glycolate metabolism at the step of glyoxylate transamination. The higher glycolate excretion observed in *lci20* (**Fig. 2b**) would then result from an inefficient conversion of glycolate into glyoxylate, which would in turn result in a decrease in the GYD1 abundance (**Fig. 3a**). Both glycolate and glyoxylate, which could not be determined in our metabolomic analysis, are intermediate metabolites whose accumulation has been shown to be toxic for plants^60,61^. Because *Chlamydomonas* is unlikely to accumulate glycolate due to the existence of an excretion pathway, we conclude that the growth impairment of *lci20* does not result from glycolate excretion per se, but rather from poisoning by non-metabolized glyoxylate. In line with our data, an *Arabidopsis* mutant deficient in the glutamate/malate transporter DiT2.1 showed photoinhibition of PSII caused by the negative feedback of non-metabolized glyoxylate on the activation state of the Calvin cycle enzymes^62,63^. Overall, LCI20 plays a vital role during the acclimation of *Chlamydomonas* from H-CO_2_to VL-CO_2_ by exporting glutamate from chloroplast toward mitochondria which is required for the function of photorespiratory pathway to efficiently recycle Rubisco oxygenation product 2-PG back to the Calvin cycle in the chloroplast.

The fact that *lci20* growth is unaffected under steady-state L-CO_2_ or VL-CO_2_ could be explained in two different ways. On the one hand, we could consider that photorespiration is only transiently active before the activation of CCM during acclimation to sub-optimal CO_2_ levels but is suppressed or strongly reduced upon CCM activation under steady-state conditions. A similar conclusion was drawn from the observation that the glycolate excretion is suppressed in air acclimated *Chlamydomonas* wild-type cells when CCM is fully induced^16^. However, we can also imagine that the arrest of glycolate excretion is caused by the upregulation of enzymes of the glycolate pathway both at transcript and protein levels. The later hypothesis is supported by the fact that the transaminase inhibitor aminooxyacetate (AOA), which blocks the photorespiratory pathway, induces a higher glycolate excretion in air-adapted *Chlamydomonas* wild-type cells with a fully activated CCM^44^. Alternatively, we could consider that during photorespiration the need for LCI20-mediated -NH2 recycling is only transient. Indeed, two -NH2 are needed for the conversion of two molecules of glyoxylate into two glycines. A -NH2 is likely supplied during the conversion of one serine into one hydroxypyruvate. Since the decarboxylation of two molecules of glycine into one serine is accompanied by the release of NH_3_, the later needs to be re-assimilated to balance the nitrogen budget. In *Chlamydomonas*, inhibition of glutamine synthase induces excretion of NH_3_ in an O_2_-dependent manner therefore showing that at least part of the NH_3_ produced by photorespiration is recycled within chloroplasts by the GS-GOGAT pathway^64^. In *Arabidopsis*, the re-assimilation of photorespiratory NH3 is mediated by the chloroplast GS/GOGAT cycle which produces glutamate in turn re-imported into mitochondria, this process being essential for photorespiration as evidenced by the lethal phenotype of both chloroplast and mitochondrial glutamate transporters^57,65,66^. In this context, the transient phenotype of *lci20* may result from the induction of an alternative pathway of NH3 reassimilation for instance through a mitochondrial isoform of the glutamate dehydrogenase^67^ which may compensate for the lack of LCI20 under steady state limiting CO_2_ (**Fig. 7**).

An hypothesis often put forward regarding the physiological role of photorespiration in microalgae is that photorespiratory metabolites could be the metabolic signal responsible for the induction of CCM during acclimation to sub-optimal CO2 levels^11,20–23,68,69^. Our data clearly establish that both the accumulation of CCM components (**Fig. 3b**) and the CCM functioning (**Fig. 2a**) are independent of O2 levels, thus ruling out this hypothesis.

This work paves the way for a more detailed understanding of photorespiration in microalgae and how it interacts with the CCM functioning. Significant amounts of glycolate have been found in the ocean, particularly during algal blooms^70–74^, suggesting that photorespiration can be active in phytoplankton despite the occurrence of CCMs in most species^9,75,76^. This apparent discrepancy could be due to the fact that CCMs from different marine species show quite different efficiencies^9,75,76^. For instance, *Nannochloropsis* has been reported to harbour a quite leaky CCM^77^. It is also possible that algal species with an efficient CCM excrete significant glycolate amounts in particular conditions, for instance when the CO_2_ concentration drops during algal blooms, or when the incident light suddenly increases. Further studies are needed to better understand how algal photorespiration contributes to the carbon footprint of oceans.

## Methods

### Growth conditions and strains

The *lci20* (LMJ.RY0402.205944), *gyd1-1* (LMJ.RY0402.206272), *gyd1-2* (LMJ.RY0402.232044), *cia5* (CC-2702), and wild-types CC4533 and CC125 were purchased from the *Chlamydomonas* resource center. Cells were cultivated in an incubation shaker (INFORS Multitron pro) maintained at 25°C, with 120 rpm shaking and constant illumination at 80 µmol m^−2^ s^−1^ supplied by fluorescent tubes delivering white light enriched in red wavelength. Cells were grown in MOPS-buffered (20 mM MOPS, pH 7.2) minimal medium (MM) exposed to various CO_2_ levels. Photorespiration was induced by transferring 2% (H) CO_2_-grown cells into 0.01% (VL) CO_2_ under atmospheric O_2_ level (21%) and can be suppressed by 2% O_2_. Growth kinetics were monitored with a Multisizer 3 Coulter counter (Beckman Coulter). For spot test, cells were grown in liquid culture with MM, harvested during active growth at around 10 µg mL^-1^ of chlorophyll and resuspended in fresh MM medium to make series of dilutions to 0.5, 1, and 2 µg mL^-1^ chlorophyll per spot. Eight-microliter drops were spotted on 1.5% of MM agar plates at pH 7.2 buffered with 20 mM MOPS and exposed to various CO_2_ and light regimes. Homogeneous light was supplied by a panel of fluorescent tubes.

### Protein extraction and immunoblot analysis

Total protein was extracted as previously described^25^. Exponentially grown cells (equivalent of 20 µg of chlorophyll) were harvested by centrifugation at 4000 *g* for 3 min at 4°C. Pellets were resuspended in 200 µL of PBS with a complete protease inhibitor EDTA-free mixture tablet (Roche) by vortexing and sonicated for 15 s 30% pulses on ice using a sonicator (product number UR-21P; TOMY). 200 μL of Novex™ Nupage™ LDS buffer 2x (Invitrogen™) containing 1x reducing agent DTT was added to the solution, and the total protein was solubilized by incubation at 37°C for 20 min. Incubated samples were subsequently centrifuged at 4000 *g* for 3 min. 10 μL (1X) of protein samples were loaded on Novex™ Nupage™ Bis tris 10% or tris-acetate 3-8% (Invitrogen™) gel, migrated 1 h at 190 V in Novex™ Nupage™ MOPS or tris-acetate (Invitrogen™) buffer according to protein molecular weight and transferred to nitrocellulose membrane using semidry transfer technique. Immuno-detection was performed using antibodies raised against LCI20, GYD1, HPR1, GCSP, HLA3, LCI1 and LCIC. Antibody raised against α-Tubulin 1/1000 (AS10 680) was used as control. Secondary anti-rabbit peroxidase-conjugated antibodies (Sigma-Aldrich; no. AQ132P) (1/10,000) were used for the detection with the G:BOX Chemi XRQ system (Syngene) using ECL detection reagents (GE Healthcare). Images were captured with a CCD camera equipped with a GeneSys Image Acquisition Software (Syngene).

### Glycolate quantification

Excreted glycolic acid was analyzed from the supernatant after centrifugation of 1 mL culture at 4000 *g* for 2 min at 4°C. The supernatant was diluted 10x in acetonitrile and analyzed using a Vanquish UHPLC/Q Exactive Plus (ThermoScientific). Glycolic acid were separated using a HILIC stationary phase (SeQuant ZIC HILIC, 100 × 2.1 mm, 3.5 µm), heated to 35°C. A binary solvent system was used, in which mobile phase A consisted of acetonitrile: water (95:5, v/v) with 5 mM ammonium acetate and mobile phase B consisted of water:acetonitrile (95:5, v/v) with 5 mM ammonium acetate. Separations were done over a 32 min period following the gradient: 0-0.5 min: 5% B, 0.5-24.5 min: 5-95% B, 24.5-26.5 min: 95% B, 26.5-26.6 min: 95-5% B, 26.6-32 min: 5% B. The flow rate was set to 0.3 mL min^-1^ and the injection volume was 5 µL. After separation, glycolic acid was directed into the ESI source of the Q Exactive Plus (Orbitrap-mass spectrometer). The ESI source was set as following: negative mode ion spray voltage at -2.5 kV, capillary temperature at 300°C, S-lens RF level at 50. Data was acquired using a targeted t-SIM method with glycolic acid in inclusion list (formula C_2_H_4_O_3_, m/z 75.00877).

### Measurement of chlorophyll fluorescence

Chlorophyll fluorescence measurements were performed using a Dual Pulse Amplitude Modulated Fluorometer (DUAL-PAM-100; Walz) equipped with a red LED source of actinic light. Samples (2 mL of cells at 10 µg mL^-1^ chlorophyll) were placed into a cuvette under constant stirring at room temperature (25°C) and dark adapted for 15 min prior to measurement. Both effective PSII quantum yield and 1-qL were calculated from the light curve. The latter was obtained by increasing red actinic light stepwise every 1 min starting from 17, 110, 190, 390, 610 and 940 μmol photons m^−2^ s^−1^, each being separated by a saturating pulse 10,000 μmol photons m^−2^ s^−1^, 600 ms duration. The effective PSII quantum yield and 1-qL were calculated as ϕPSII= (*F*m’-*F*s)/*F*m’ and qL= ((*F*m’-*F*s)/(*F*m’-*F*0))*(*F*0/*F*s) respectively with *F*m’ the fluorescence value after saturating pulses, *F*s the stationary fluorescence at each actinic light and *F*0 the fluorescence value in the dark^78^.

### Metabolomics analysis

Cells were grown to exponential phase under photoautotrophic H-CO_2_ conditions in the flask with 80 μmol photons m^−2^ s^−1^. For L-CO_2_ and VL-CO_2_ conditions, H-CO_2_ grown cells were transferred to either L-CO_2_ or VL-CO_2_ levels for 20 h prior to sampling. About 60 million cell suspensions were injected into a -70°C cold quenching solution composed of 70% methanol in water using a thermoblock above dry ice to obtain a final concentration of 35% methanol. Centrifuge tubes containing the cell suspensions were cooled in a pre-chilled cooling box to keep the sample temperature below -20°C. Cell pellets were collected by centrifugation at 4000 *g* for 2 min at -10°C. The supernatant was decanted and residual liquid together with cells carefully transferred to a 2-mL Eppendorf tube and re-centrifuged 13000 *g* for 1 min at -10°C. The pellet was flash-frozen in liquid nitrogen and lyophilized at -50°C. The extraction and derivation of the metabolites were performed as previously^79^. The analytic and quantification methods were exactly as reported in^36^. Data is reported following recently updated metabolomics standards^80^.

### Statistics

All statistical tests used are noted in figure legends. One-way ANOVA using GraphPad Prism (GraphPad Software) was used to perform statistical analysis. The P-values were computed by One-way ANOVA test (uncorrected *p* values). Statistical significance (α = 0.05) according to *p* values is indicated by asterisk (* for *p* ≤ 0.05; ** for *p* ≤ 0.01; *** for *p* ≤ 0.001 and **** for *p* ≤ 0.0001).

### Data availability

Genes studied in this article can be found on Phytozome (v6.1 genome: https://phytozome-next.jgi.doe.gov/) under the loci Cre06.g260450 (LCI20), Cre06.g295450 (HPR1), Cre06.g288700 (GYD1), Cre12.g534800 (GCSP), Cre03.g162800 (LCI1), Cre02.g097800 (HLA3), Cre06.g307500 (LCIC) and Cre02.g096300 (CIA5). All antibodies used are available upon request to corresponding authors. All source data are provided with this paper as a Source Data File.

All the other methods are reported in the Supplemental Information file.

## Acknowledgment

O.D. thanks the French Atomic Energy and Alternative Energy Commission (CEA) for a PhD scholarship. We thank Shiyan Zheng for assistance in performing the genetic complementation experiment; Gaurav Kumar for technical assistance during microscopic observation; Stephanie Blangy for trials on generating antibodies against LCI20; Franziska Kuhnert for trials on the transport assays. We thank the ZoOM Microscopy facility (CEA Cadarache) and the University of York Biosciences Technology Facility for confocal microscopy access. This work is financially supported by the “L’économie Circular du Carbone” program of the CEA (CO2storage), the ANR project “AlgalCCM” (n°ANR-366 22-CE44-0023-01), the France 2030 initiative project “CO2_CMPhi” (ANR-23-PEXF-0002), and the Region Sud (“AlgalCO2” project). We acknowledge the European Union Regional Developing Fund (ERDF), the Région Provence Alpes Côte d’Azur, the French Ministry of Research and the CEA for funding the HelioBiotec platform. C.M. is grateful to the INRAE MIGALE bioinformatics facility (MIGALE, INRAE, 2020) for providing computing resources. AKL was supported by the Labex Saclay Plant Sciences-SPS (ANR-17-EUR-0007), the platform of Biophysics of the I2BC supported by the French Infrastructure for Integrated Structural Biology (FRISBI; grant number ANR-10-INSB-05). A.P.M.W. acknowledges funding by the European Union’s H2020 research and innovation and the Horizon programs (grants GAIN4CROPS, GA No. 862087 and BEST-CROP, GA No. 101082091) and the Deutsche Forschungsgemeinschaft (Cluster of Excellence for Plant Sciences (CEPLAS) under Germany’s Excellence Strategy EXC-2048/1 under project ID 390686111). S.A, and A.R.F. acknowledge the European Union’s Horizon 2020 research and innovation program, project PlantaSYST (SGA-CSA No. 739582 under FPA No. 664620) and the BG05M2OP001-1.003-001-C01 project, financed by the European Regional Development Fund through the Bulgarian ‘Science and Education for Smart Growth’ Operational Program. Both authors acknowledge the support by the Max Planck Society. A.B. acknowledges support from the Carnegie Institution for Science.

## Author Contributions

Y.L-B., G.P., O.D. conceived the study. Y.L-B. and G.P. supervised the work. O.D. performed most of the experiments. A.B., A.K.L. and G.P. supervised photosynthesis measurement performed by O.D.. P.A., M.B. and O.D. carried out immunoblots. O.D. and V.E. isolated and characterized *gyd1* mutants with contribution from A.M.. C.M. performed phylogenetic analysis. M.B., F.V., and O.D. performed genetic complementation of *lci20* mutant. S.A. and A.R.F. performed metabolomics. O.D., P.-C.N. and B.L. performed glycolate analysis. O.D., G.P. and Y.L-B. drafted the manuscript with contributions from L.C.M.M., A.B., A.K.L., A.R.F., and A.P.W..

## Competing interests

there is no competing interest.

## Notes

### Competing Interest Statement

The authors have declared no competing interest.

