## Supplementary information for "Joint operation of the CO_2_ concentrating mechanism and photorespiration in green algae during acclimation to limiting CO_2_"

**This PDF file includes:**

Supplementary Methods
Supplementary figures 1 to 6
Supplementary References

### Supplementary Methods

**Genetic complementation and *Chlamydomonas* transformation.** For the construction of *pPSAD::LCI20:tPSAD*, full-length genomic sequence of *LCI20* was obtained by PCR using the high fidelity KOD Hot Start DNA Polymerase (Merck Millipore) from *Chlamydomonas* genomic DNA (extracted as previously described) using *LCI20* specific primers LCI20-fwd-ATG (ATGAGTGCACTTCTGGCTAG) and LCI20-rev-TAA (TTACCACCAGCCCAGCAGCT) flanked by the restriction site of *Bbs*I enzyme. The PCR products were cloned under the control of the *PSAD* promoter and terminator. For transformation, the fragment harboring hygromycin resistance gene in addition to *pPSAD::LCI20:tPSAD* from the digested plasmid was incorporated into the genome of exponentially grown *Chlamydomonas* cells by electroporation and spread on TAP hygromycin (15 mg $L^{-1}$) agar plates. Transformants were screened by PCR using *LCI20* specific forward and reverse (TCATCGTGGTGTCGTTCTTCTTCGC and TGCGCTCTCCCAGGCCCGTCTTCTC) primers respectively. *RACK1* was amplified as a control gene using forward (GAGTCCAACTACGGCTACGCC) and reverse (CTCGCCAATGGTGTACTTGCAC) primers. The growth of hygromycin-resistant colonies was assessed by spot test on agar plate under VL-$CO_2$ conditions to screen for positive complemented lines.

***Chlamydomonas* confocal microscopy.** LCI20-Venus transgenic strain (CSI_FC1G02) harboring the pLM005-Cre06.g260450-Venus-3xFLAG construct was ordered from the *Chlamydomonas* center[1,2]. CC4533 and LCI20-Venus cells were grown mixotrophically in TAP media prior to imaging. Cells were mounted on 18-well chamber slides and overlayed with 1.5% low melting point agarose made with TP-medium. Images were obtained with a LSM880 (Zeiss) equipped with an Airyscan module using a 63× objective. Laser excitation and Emission setting for each channel used were the following: Venus (Excitation: 514 nm; Emission 525 – 500 nm) and Chlorophyll (Excitation: 633 nm; Emission 670 – 700 nm).

**Generation of antibodies.** Antipeptide antibodies were made by immunizing two rabbits with synthetic peptides against LCI20 (DAPSSQNGVHHDPVPEC and CDDSRKKMGSYLIQSQ), GYD1 (GEVNRILAAHQKKNKL), HPR1 (SNYAVGYNNVKVDEATKRC), GCSP (SAIARGKKPKFLVSSKC), HLA3 (RKMAEDFWSTRSAQGRNQ), LCI1 (DAEESHAMPNVHVTSDGATKV) and LCIC peptides as described[3]. Synthesis of the peptides, rabbit immunization, and purification of antibodies were performed by Proteogenix SAS (Schiltigheim). Specificities of the antibodies were then tested on total proteins extracted from whole-cell *Chlamydomonas* parental lines (CC125 and CC4533) and the *cia5* mutant.

**Phylogenetic analysis.** A phylogenetic tree of the dicarboxylate transporter family was built from reviewed sequences of the InterPro entry IPR030676 / CitT-rel (December 2023) and were completed with a selection of homologous sequences from the *Viridiplantae*, including *Chlorophyta*, and malate transporters from *Eubacteria*. The multiple sequence alignment was performed with MAFFT v7.49[4] using the L-INS-i strategy, and filtered for saturation with BMGE v1.12[5] using the BLOSUM62 matrix and removing sites with > 50% gaps without entropy-based trimming, to get a final alignment containing 462 sites and 66 sequences. The tree was built using the Maximum-Likelihood (ML) algorithm with IQ-TREE[6] and the Q.LG+F+I+G4 substitution model selected by ModelFinder[7] with the Bayesian Information Criterion (BIC) and restricting model selection to the cpREV, WAG, Q.plant, Q.pfam and Q.LG matrices

(**Supplementary Fig. 2**). The statistical support of the branches was estimated using a (UFBoot) approximation[8] implemented in IQ-TREE with 1000 replicates. A second tree was built from a reduced dataset of 21 sequences inspired from[9,10] (**Supplementary Fig. 1c**) using the same approach.

### Supplementary Figures

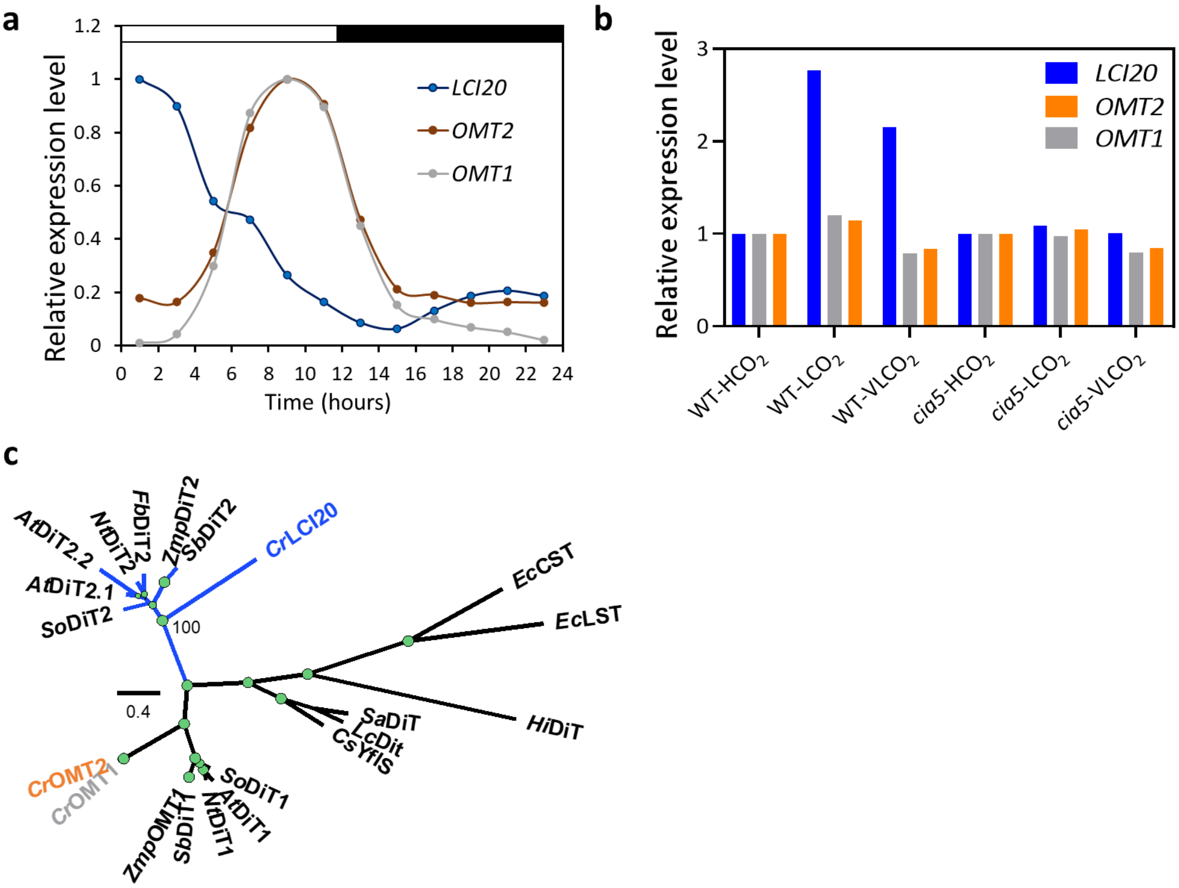


**Supplementary Fig. 1. *LCI20* putatively encodes a malate transporter expressed under low $CO_2$ or at the onset of illumination during a day-night cycle.** (**a**) Expression level of *LCI20, OMT1* and *OMT2* genes in *Chlamydomonas* WT during a day/night cycle. Data were obtained from[11]. (**b**) Expression level of *LCI20, OMT1* and *OMT2* genes in *Chlamydomonas* WT acclimated to different $CO_2$ levels. Data were obtained from[12]. (**c**) Unrooted Maximum Likelihood tree of the dicarboxylate transporter family in plants, *Chlamydomonas reinhardtii* and *Eubacteria* showing the affiliation of the *LCI20* gene to the Dit2 group. The first two letters of the acronyms indicate the species (At, *Arabidopsis thaliana*; Cr, *Chlamydomonas reinhardtii*; Cs, *Clostridium saccharobutylicum*; Ec, *Escherichia coli*; Fb, *Flaveria bidentis;* Hi, *Haemophilus influenzae*; Lc, *Liquorilactobacillus cacaonum*; Nt, *Nicotiana tabacum*; Sa, *Staphylococcus aureus*; So, *Spinacia oleracea*; Zm, *Zea mays*). The three following letters indicate the group of transporters (DiT, dicarboxylate transporter; TST, L-tartrate/succinate transporter; CST, citrate/succinate transporter; YlfS, 2-oxoglutarate/malate transporter). The tree was drawn to scale. The scale bar represents the number of substitutions per site. Tree topology was tested using an ultrafast bootstrap approximation approach with 1000 replicates. Gray circles represent bootstrap values and are drawn to scale. Dotted branches represent bacterial lineages. A full version of this tree with primary accession numbers is given in **Supplementary Fig. 2**.

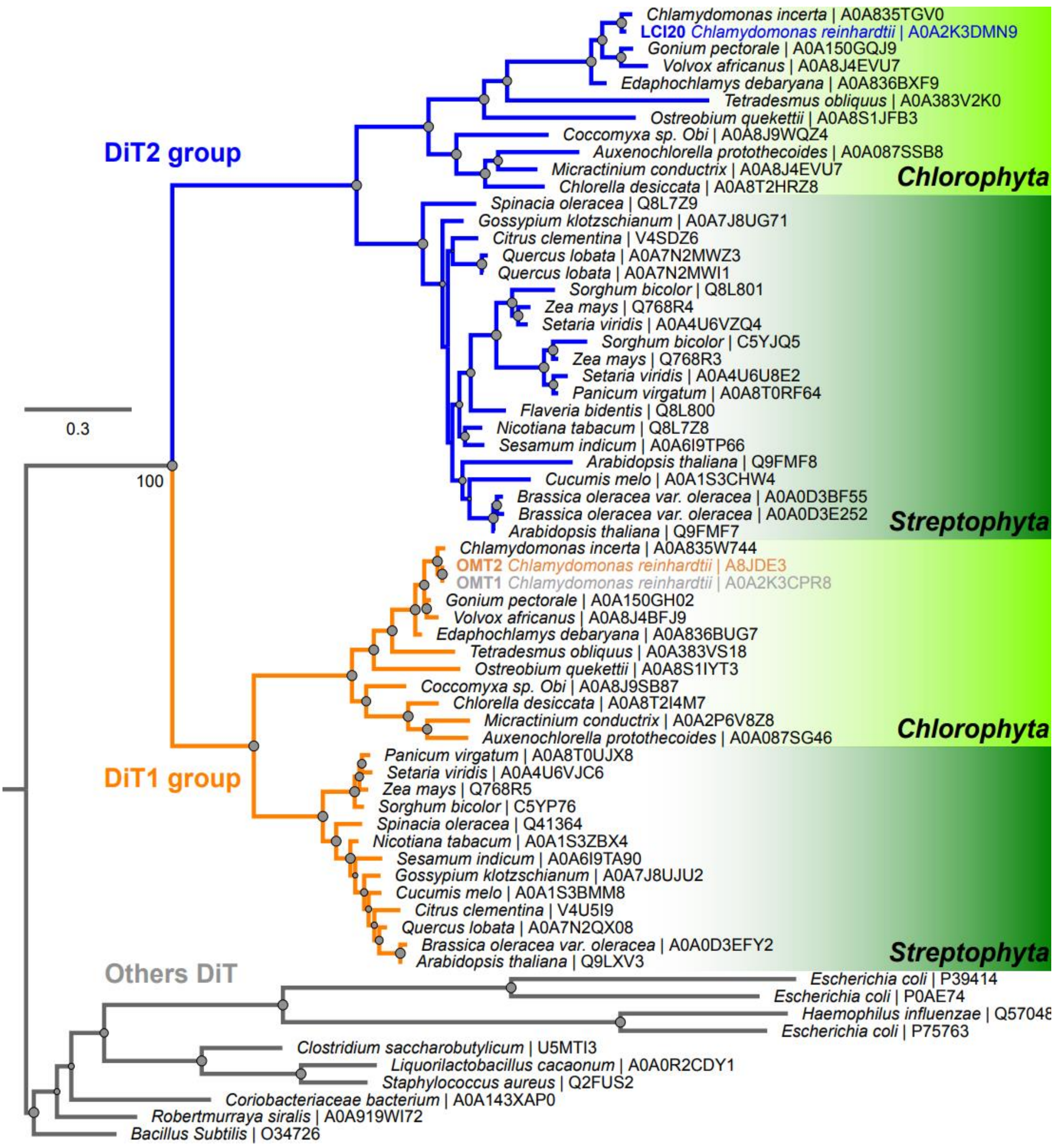


**Supplementary Fig. 2. Maximum likelihood tree of the dicarboxylate transporter family (Citrate carrier CitT-related / IPR030676).** The tree was rooted with *Eubacteria* sequences. Each sequence is associated to its primary accession number in the public database UniProtKB (https://www.uniprot.org). The tree was drawn to scale. The scale bar represents the number of substitutions per site. Tree topology was tested using an ultrafast bootstrap approximation approach with 1000 replicates. Gray circles represent bootstrap values and are drawn to scale.

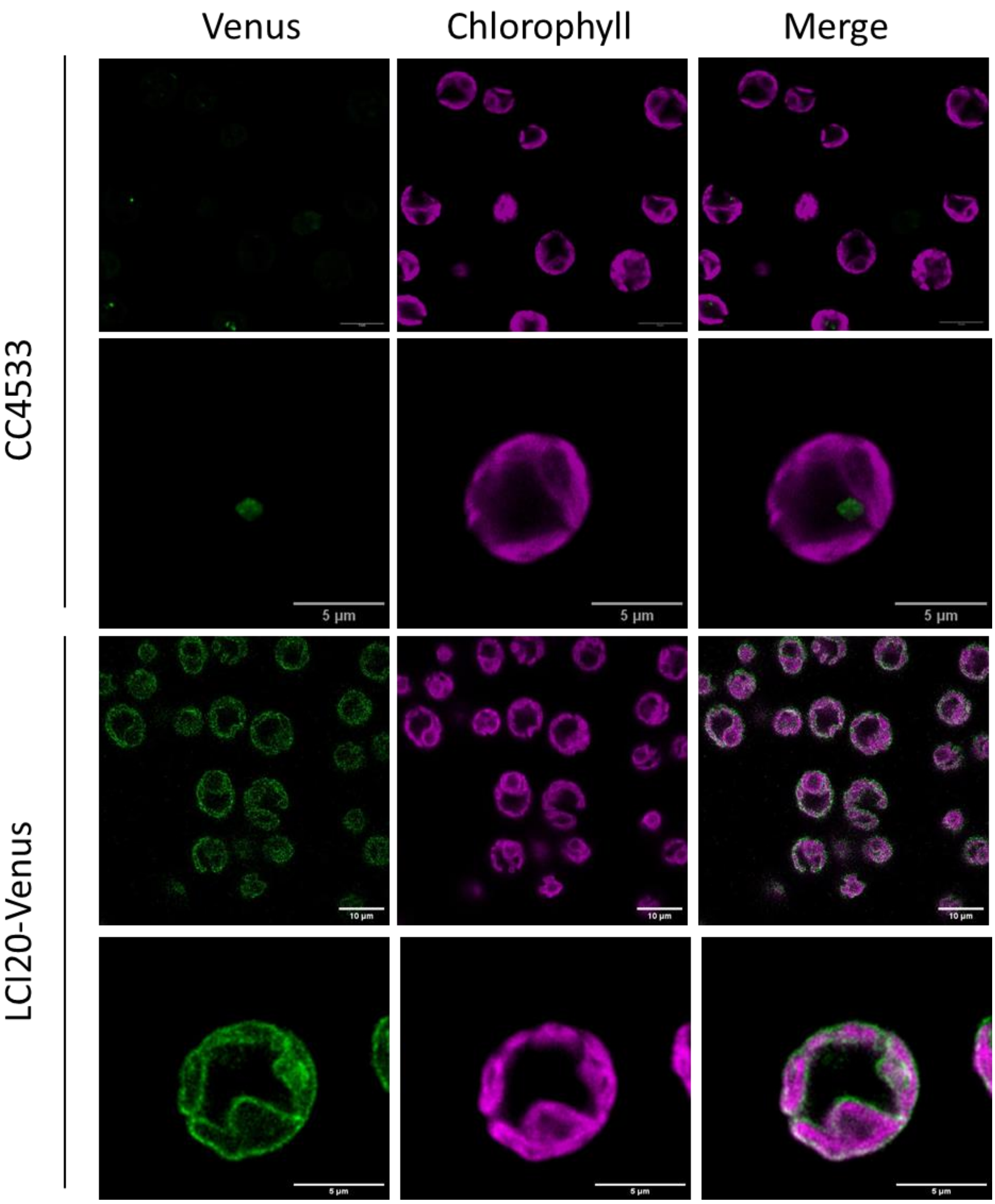


**Supplementary Fig. 3. Subcellular localization of LCI20 protein fused with the Venus fluorescent reporter.** False colours were used to represent Venus (green) and chlorophyll (magenta) fluorescence signal. Multiple cells (top panel) and individual cell (bottom panel) are shown from CC4533 and LCI20-Venus strains. Merge represents overlayed channel of Venus and chlorophyll autofluorescence. Scale bar is 5 or 10 µm as indicated.

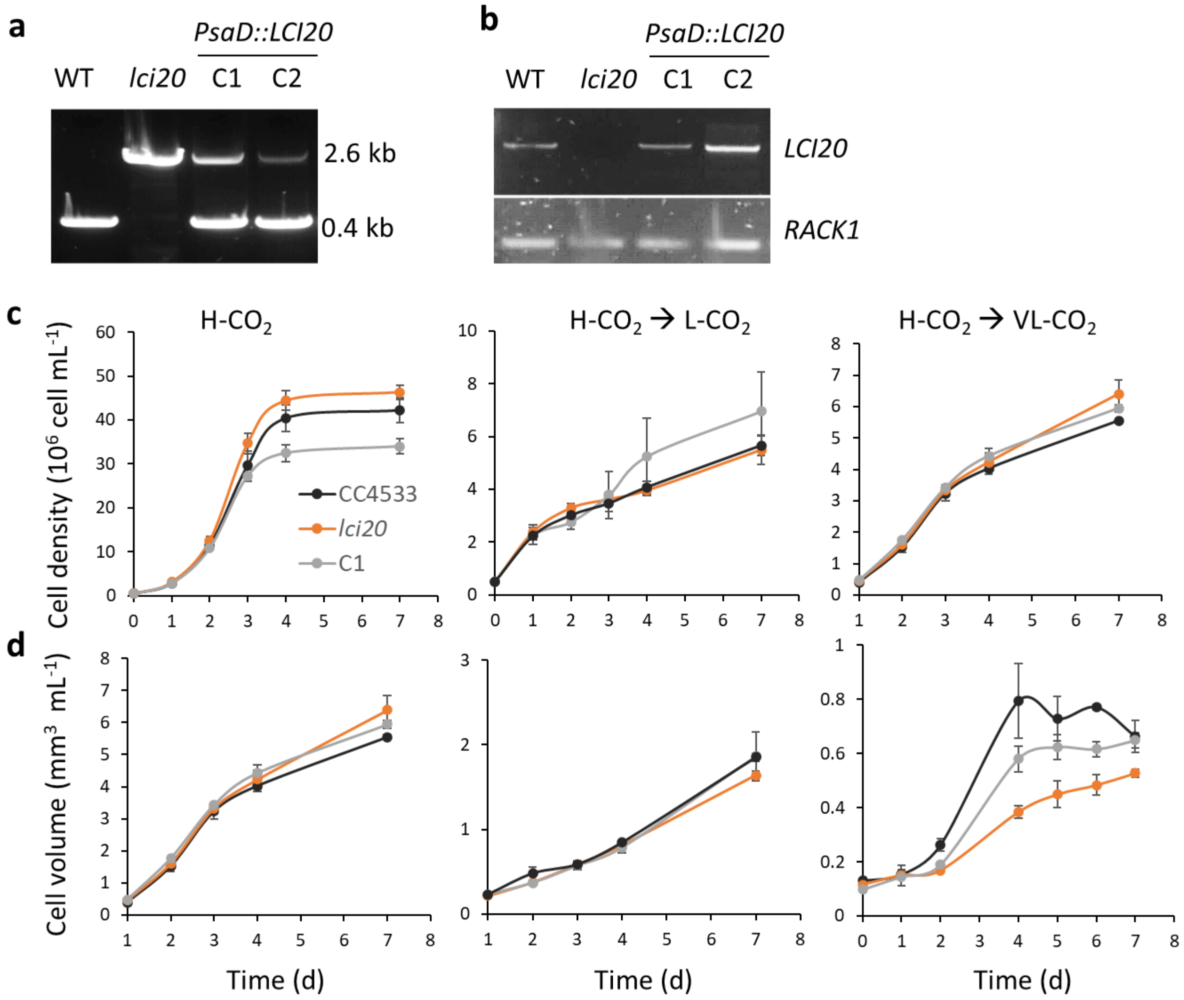


**Supplementary Fig. 4. Genotyping *lci20* and of two complemented strains and growth in liquid photoautotrophic cultures under various $CO_2$ concentrations.** (**a**) Genotyping and validation of genetic complementation of *lci20* insertional mutant by PCR. The illustration shows PCR amplification of the *LCI20* gene using primers flanking the paromomycin resistance gene insertion, resulting in a larger PCR product in *lci20* and the complemented lines showing both the endogenous *LCI20* and the transgenic full length *LCI20*. (**b**) RT-PCR showing the absence of *LCI20* transcript in the *lci20* mutant and the restauration of *LCI20* expression in the complemented lines. *RACK1* cDNA was amplified as a control gene. (**c**) Photoautotrophic growth in liquid cultures of *lci20*, its wild-type control and a complemented line following the cell number over 7 days under H-$CO_2$, during acclimation to L-$CO_2$, and VL-$CO_2$ 21% $O_2$. (**d**) Same as (**c**) but growth was followed by measuring the total cellular volume. Photoautotrophic H-$CO_2$ grown cells were used to inoculate fresh MM growth media at 0.5 million cells $mL^{-1}$ cell density at day zero. Light intensity was 80 μmol photons $m^{-2}$ $s^{-1}$ throughout the experiment. Data are means of three biological replicates for each strain ± SD.

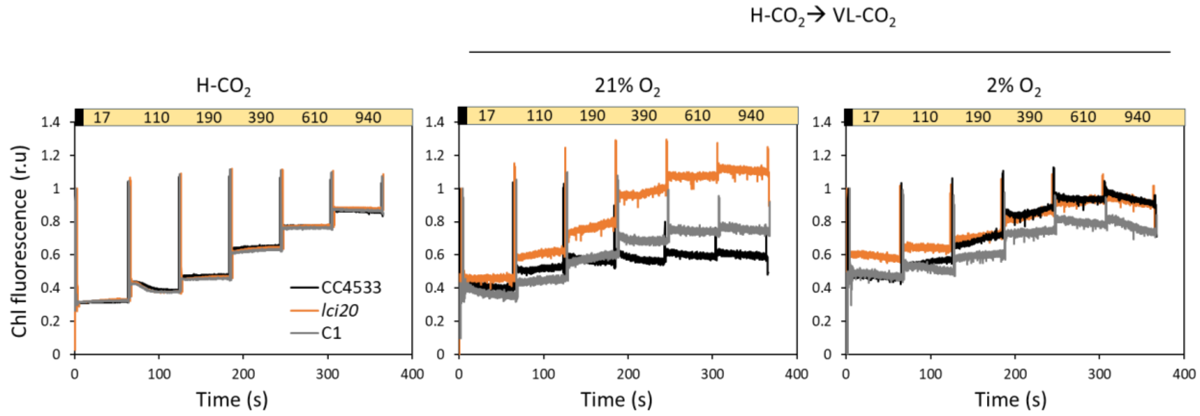


**Supplementary Fig. 5. Representative chlorophyll fluorescence traces of *lci20*, its wild-type control and one complemented line recorded under various $CO_2$ levels.** Chlorophyll fluorescence measurements were carried out using actinic red light of stepwise increasing intensity: 0, 17, 110, 190, 390, 610 and 940 µmol photons $m^{-2}$ $s^{-1}$ respectively. The black and yellow boxes represent the dark and light phases respectively. Saturating flashes, indicated by vertical lines, were supplied every 60s. Data are normalized on initial F*m* measurements and traces are shifted few seconds to allow clarity. The initial F*m* and $F_0$ was determined after a 15 min dark adaptation. Chlorophyll fluorescence traces such as the ones shown here were used to calculate data shown in Fig. 4 as described in methods section.

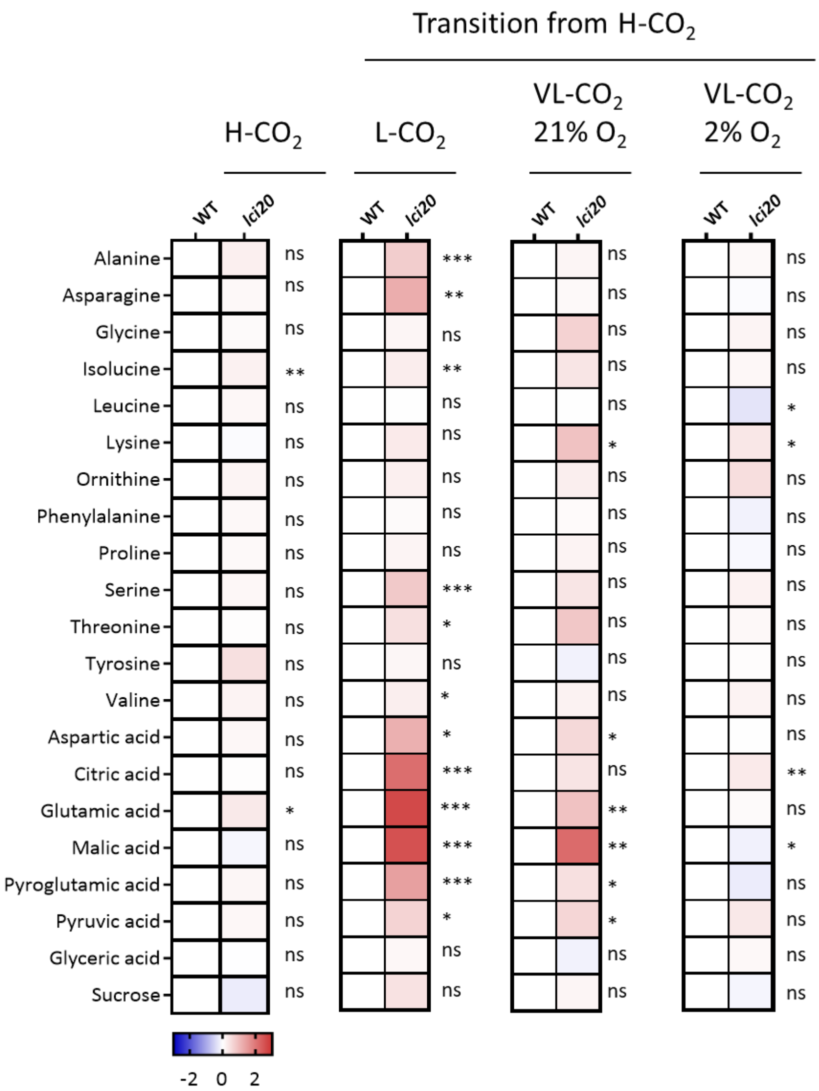


**Supplementary Fig. 6. Metabolomics analysis of *lci20* and its wild-type control acclimated to H-$CO_2$ and during acclimation to L- or VL-$CO_2$ conditions.** Heatmap showing fold-changes of metabolites in the *lci20* as compared to the WT under H-$CO_2$, and during acclimation to L-$CO_2$, VL-$CO_2$ 21% $O_2$ and VL-$CO_2$ 2% $O_2$ for 20 h. Shown represents the mean of eight biological replicates for each strain. Asterisks represent statistically significant differences compared to WT CC4533 (* $p \leq 0.05$, ** $p \leq 0.01$ and *** $p \leq 0.001$) using one-way ANOVA.
